# ntHits: *de novo* repeat identification of genomics data using a streaming approach

**DOI:** 10.1101/2020.11.02.365809

**Authors:** Hamid Mohamadi, Justin Chu, Lauren Coombe, Rene Warren, Inanc Birol

## Abstract

**Motivation:** Repeat elements such as satellites, transposons, high number of gene copies, and segmental duplications are abundant in eukaryotic genomes. They often induce many local alignments, complicating sequence assembly and comparisons between genomes and analysis of large-scale duplications and rearrangements. Hence, identification and classification of repeats is a fundamental step in many genomics applications and their downstream analysis tools.

**Results:** In this work, we present an efficient streaming algorithm and software tool, ntHits, for *de novo* repeat identification based on the statistical analysis of the *k*-mer content profile of large-scale DNA sequencing data. In the proposed algorithm, we first obtain the *k*-mer coverage histograms of input datasets using the ntCard algorithm, an efficient streaming algorithm for estimating the *k*-mer coverage histograms. From the obtained *k*-mer coverage histogram, the repetitive *k*-mers would present a long tail to the distribution of *k*-mer coverage profile. Experimental results show that ntHits can efficiently and accurately identify the repeat content in large-scale DNA sequencing data. For example, ntHits accurately identifies the repeat *k*-mers in the white spruce sequencing data set with 96× sequencing coverage in about 12 hours and using less than 150GB of memory, while using the exact methods for reporting the repeated *k*-mers takes several days and terabytes of memory and disk space.

**Availability:** ntHits is written in C++ and is released under the MIT License. It is freely available at https://github.com/bcgsc/ntHits.

## 1 Introduction

Repeat elements are widespread among among eukaryotes and form a major portion of their genomes. They are usually classified on the basis of their sequence characteristics and of how they are formed. One category consists of tandem repeats, and includes any sequences found in consecutive copies along a DNA strand. Several different categories of tandem repeats have been defined, depending on the number of repeats and on the size of the repeated units. This group includes microsatellites or simple sequence repeats and minisatellites. Another category, on which this review will mainly focus, is constituted by elements that are found dispersed across the whole genome, and which consists mainly of TEs. TEs can be classified according to the intermediate they use to move.

Repeat elements such as satellites, transposons, high number of gene copies, and segmental duplications are abundant in eukaryotic genomes. They often induce many local alignments, complicating sequence assembly and comparisons between genomes and analysis of large-scale duplications and rearrangements. Hence, identification and classification of repeats is a fundamental step in many genomics applications and their downstream analysis tools. Currently most repeat analysis tools are exact methods that rely on reference genomes and are based on costly similarity searches. On the other hand, *de novo* computational identification of such elements is a very computationally challenging problem. The existing tools for this purpose need considerable computational resources in terms of memory, disk space, and runtime requirements for processing and identifying repeats in large sets of DNA sequences.

In this paper we introduce a streaming method to identify repeat *k*-mers in large-scale NGS data. Streaming algorithms are algorithms for processing data that are too large to be stored in available memory, but can be examined online, typically in a single pass. There has been a growing interest in streaming algorithms in a wide range of applications, in different domains dealing with massive amounts of data. Examples include, analysis of network traffic, database transactions, sensor networks, and satellite data feeds (Cormode and Garofalakis, 2005; Cormode and Muthukrishnan, 2005; Indyk and Woodruff, 2005). In bioinformatics they have been used in some applications for exploring genome characteristics and quality of the sequencing data (Chikhi and Medvedev, 2014; Melsted and Halldorsson, 2014; Simpson, 2014) Our proposed method will be based on statistical analysis of *k*-mer depth frequency histogram obtained using the ntCard algorithm. From the computed *k*-mer frequency histograms for a given dataset, the proposed method applies tries to fit a mixture model to deconvolute different distribution for the combined frequency histogram. After identifying distributions for error, heterozygous, homozygous, and repeat *k*-mers, the proposed approach estimate the threshold for repeat *k*-mers. It then process the input data in the second pass to extract the *k*-mers appearing more than the estimated repeat threshold value.

## 2 Method

In the proposed algorithm, we first obtain the *k*-mer coverage histograms of input datasets using the ntCard algorithm (Mohamadi *et al*., 2017). ntCard is an efficient streaming algorithm for estimating the *k*-mer coverage histograms from raw NGS. It works by first hashing the *k*-mers in read streams, which it samples to build a reduced multiplicity table. After calculating the multiplicity table for sampled *k*-mers, it uses this table to infer the population histogram through a statistical model. ntCard utilizes the ntHash algorithm (Mohamadi *et al*., 2016) to efficiently compute the canonical hash values for all *k*-mers in DNA sequences. ntHash is a recursive, or rolling, hash function in which the hash value for the next *k*-mer in an input sequence of length *l* (*l* ≥ *k*) is derived from the hash value of the previous *k*-mer.

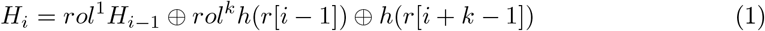

This calculation is initiated for the first *k*-mer in the sequence using the base function

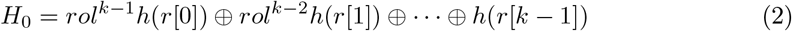

Where ⊕ is the bitwise exclusive or operation, *rol* is cyclic binary left rotation, and *h* is a seed table mapping the nucleotide letters to a pre-designed 64-bit random integers.

From the obtained *k*-mer coverage histogram, the repetitive *k*-mers would present a long tail to the distribution of *k*-mer coverage profile. On the other hand, experimental noise from base calling errors would typically result in *k*-mers with low frequencies, if the experimental noise were not systematic. Thus, the probability of observing the erroneous *k*-mers should follow an exponential distribution. To make the distribution fitting more robust, we first exclude the erroneous *k*-mers that are likely caused by sequencing errors by fitting an exponential distribution to the *k*-mer coverage histogram and then identifying a threshold for erroneous *k*-mers denoted by λ_*e*_. After excluding the effect of erroneous *k*-mers frorm the histogram, ntHits identifies the homozygous *k*-mers (*k*-mers common in both parental alleles) rate, denoted by λ, which is usually the maximum peak in the histogram. The *k*-mers that are repeated in the input data occur at rate λ_*r*_ = *r* × λ, *r* ≥ 2. After identifying the error rate, λ_*e*_, and repeat rate, λ_*r*_, parameters, ntHits streams through the input datasets and then filters out non-repetitive *k*-mers using a counting Bloom filter data structure. The counting Bloom filter will count up to the repeat threshold λ_*r*_ and the *k*-mers that appear more than the repeat threshold will be passed to the next stage and stored in a hash table with their counts. The estimated number of repeated elements that will be stored in the hash table will also be identified from the *k*-mer coverage histogram obtained by the ntCard algorithm.

### 2.1 Constructing *k*-mer frequency histograms using ntCard

Figure 1 shows the ntCard workflow to construct the k-met depth frequency histogram form raw NGS reads. As input we are given a set of raw NGS data. ntCard uses ntHash algorithm to compute the 64-bit hashes for all *k*-mers in input data. After computing hashes, they are divided in three segments. ntCard uses the *s* left bits to sample from input data that results in a sampled data of size 1*/*2^*s*^. After sampling, *r* right bits of hashes are used to build a *k*-mer count table. Next, from the count table, frequencies and relative frequencies are computed. After computing sample statistics, the population relative frequencies are constructed from the sample relative frequencies using a statistical model. Finally, the estimated population frequencies are computed from relative frequencies.

**Figure 1:**
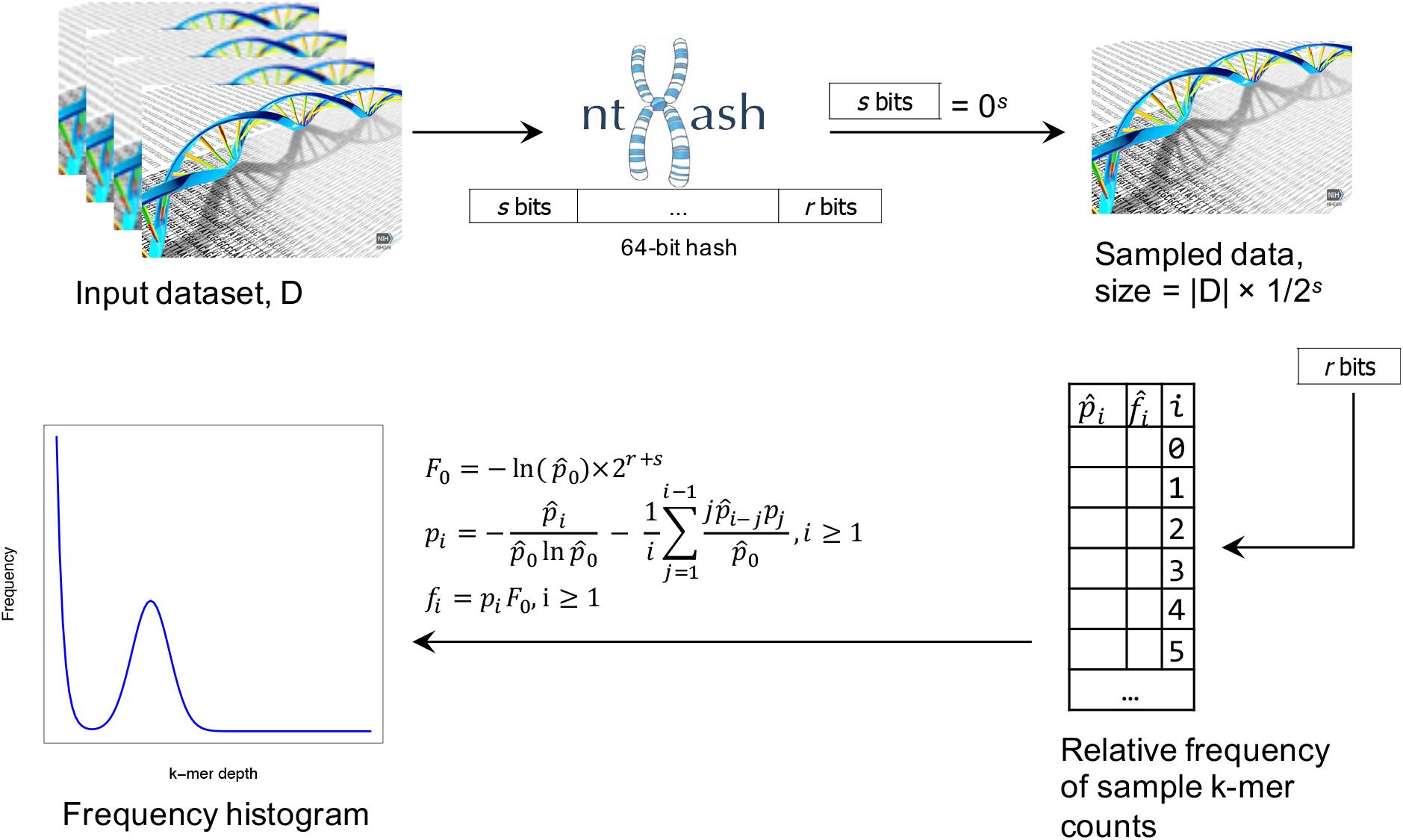
Workflow to construct *k*-mer frequency histogram using ntCard. The process starts by sampling from input data. Then the relative frequency of sample *k*-mer counts is computed. Finally, the population relative frequencies are constructed from the sample relative frequencies using a statistical model.

An example of *k*-mer frequency histogram from ntCard is shown in Figure 2.a.. As we see here, the *k*-mer hist is a combination of different distributions. The erroneous *k*-mers shown in red follow an exponential distribution. The heterozygous and homozygous *k*-mers shown in yellow and blue follow a bell-shaped distribution. Repeat *k*-mers would present a long tail to the distribution of *k*-mer histograms. In addition, the ploidy of the organism, and sequence variants in haplotypes also impact the *k*-mer histogram. Since heterozygous *k*-mers and erroneous *k*-mers have highly overlapping distributions, it is hard to determine which *k*-mers might represent haplotypic variations.

**Figure 2:**
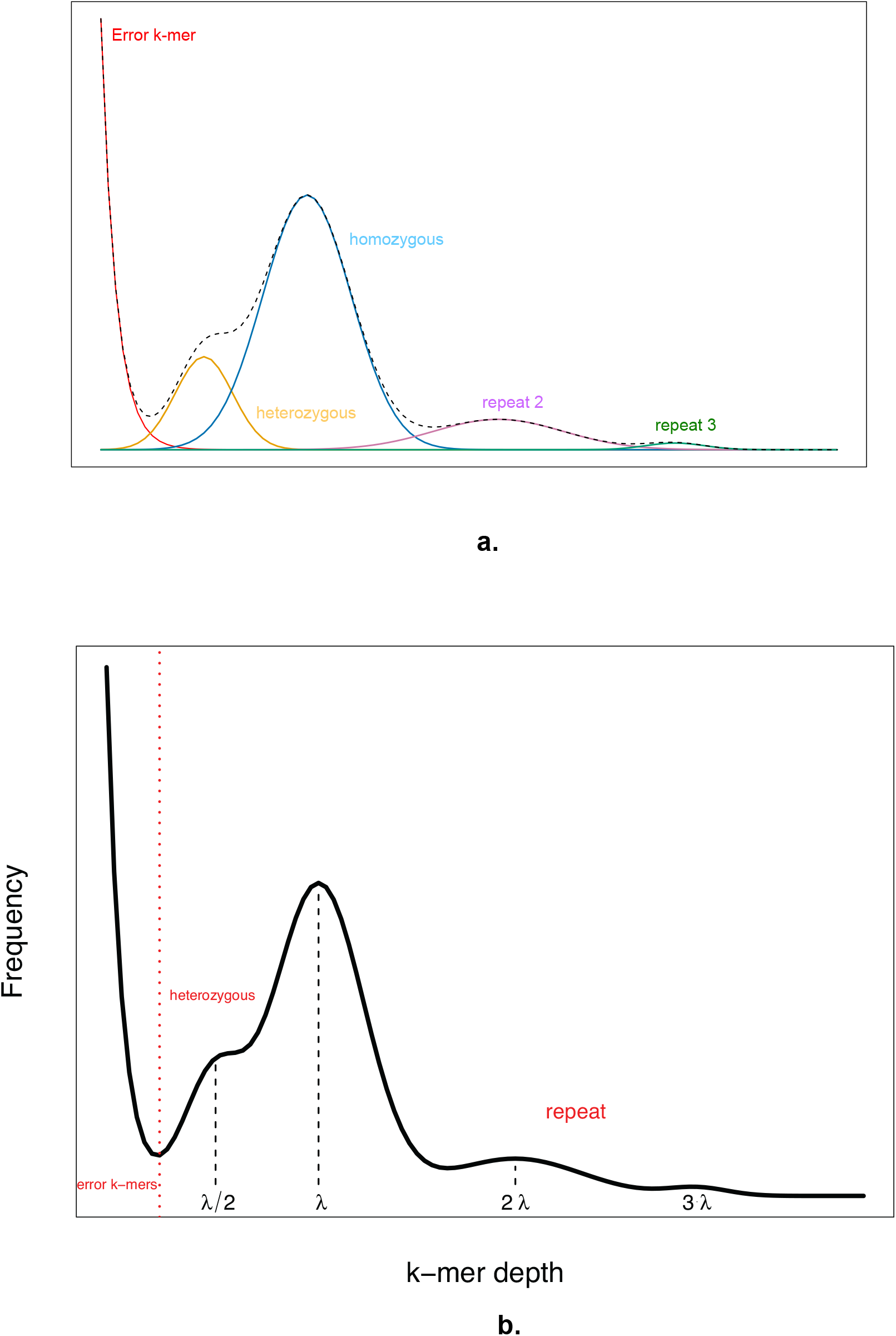
The *k*-mer frequency histogram as a mixture of different distributions.

Figure 2.b. represents a summarized version of the *k*-mer depth frequency histogram. We use a mixture of negative binomial model terms rather than Poisson terms because real sequencing data is often over-dispersed compared to a Poisson distribution and the size term can independently control the variance. The heterozygous *k*-mer rate has been denoted by a rate of λ*/*2. The homozygous *k*-mer rate has been denoted by λ. 2λ and 3λ are the rates for *k*-mers repeated twice, three times, and so on and so forth. Model fitting has been performed using a non-linear least squares estimate implemented by the nls function in R.

## 3 Results

### 3.1 Implementation details

In the implementation of ntHits two Bloom filters were used. The first Bloom filter keeps the track of distinct *k*-mers in the whole input dataset. The *k*-mers with count greater than 1 will pass through this step. The second Bloom filter is a counting Bloom filter that will count up to the repeat threshold λ_*r*_ so the *k*-mers that appear more than the repeat threshold will be passed to the next stage and stored in a Repeat Bloom filter/hash table. If input reads or sequences contain ambiguous bases, or characters other than {*A,C,G,T*}, the proposed tool ignores them in the hashing stage. This is performed as a functionality of ntHash algorithm.

ntHits is written in C++ and parallelized using OpenMP for multi-threaded computing on a single computing node. As input, it gets the set of sequences in FASTA, FASTQ, SAM, and BAM formats. The input sequences can also be in compressed formats such as .gz and .bz formats. The proposed method is distributed under the MIT License. Documentation and source code are freely available at https://github.com/bcgsc/ntHits.

### 3.2 Experimental setup

To evaluate the performance and accuracy of our proposed approach, we downloaded the following publicly available sequencing data.

- The Genome in a Bottle (GIAB) project (Zook *et al*., 2016) sequenced seven individuals using a large variety of sequencing technologies. We downloaded 2×250 bp paired-end Illumina whole genome shotgun sequencing data for the Ashkenazi mother (HG004).
- To represent a larger problem, we used the white spruce (*Picea glauca*) genome sequencing data that represents the genotype PG29 (Warren *et al*., 2015) (accession number: ALWZ0100000000 and PID: PRJNA83435).

The information of each dataset including the number of sequences, size of sequences, total number of bases, and total input size of datasets is presented in Table 1.

**Table 1:**
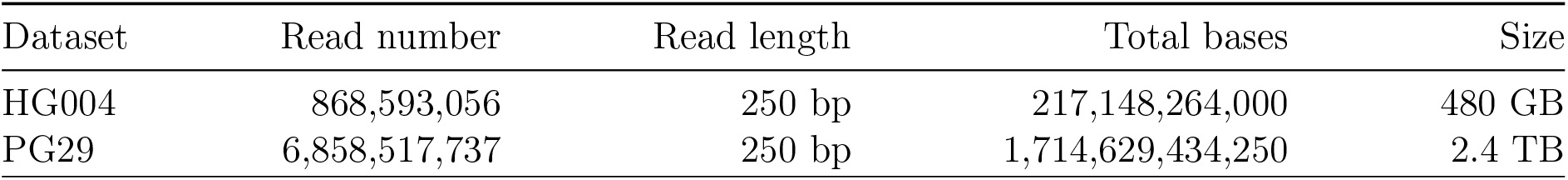
Dataset specification.

| Dataset | Read number | Read length | Total bases | Size |
| --- | --- | --- | --- | --- |
| HG004 | 868,593,056 | 250 bp | 217,148,264,000 | 480 GB |
| PG29 | 6,858,517,737 | 250 bp | 1,714,629,434,250 | 2.4 TB |

Table 2 shows the results for running ntHits on HG004 and PG29 datasets. Here, the exact results for identifying repeated *k*-mers were obtained from DSK tool (Rizk *et al*., 2013). As we see here, ntHits is able to identify repeated *k*-mers with high accuracy rate for HG004 and PG29 data with fpr rate of 0.0001 and 0.0006 respectively. The runtime and memory usage for ntHits when running on HG004 data were 2h:17m and 20GB while these values were 14h:03m and 250GB for PG29 dataset. Using exact methods for this purpose will take days and tera bytes of memory while ntHits identifies the repeated *k*-mers in large-scale sequencing datasets efficiently due to its streaming nature. ntHits has been used in the ntEdit (Warren *et al*., 2020) tool pipeline for filtering out repeated k-mers and we also expect it provides utility in efficiently characterizing certain properties of large read sets, helping quality control pipelines and de novo sequencing projects.

**Table 2:** Accuracy of the proposed algorithm in estimating repeated *k*-mers for HG004 reads. The exact results obtained from DSK *k*-mer counting tool.

|  |  | Exact<br>HG004 |  |
| --- | --- | --- | --- |
|  |  | repeat | non-repeat |
| Predicted | repeat | 54,583,915 | 5,554 (fpr = 0.0001) |
|  | non-repeat | 0 | 19,798,490,149 |
|  |  | PG29 |  |
| Predicted | repeat | 5,259,022,647 | 3,400,484 (fpr = 0.0006) |
|  | non-repeat | 0 | 55,268,063,627 |

